# Studying 3D cell cultures in a microfluidic droplet array under multiple time-resolved conditions

**DOI:** 10.1101/407759

**Authors:** Raphaël F.-X. Tomasi, Sébastien Sart, Tiphaine Champetier, Charles N. Baroud

## Abstract

The relevance of traditional cell cultures to cellular behavior *in vivo* is limited, since the two-dimensional (2D) format does not appropriately reproduce the microenvironment that regulates cell functions. In this context, spheroids are an appealing 3D cell culture format to complement standard techniques, by combining a high level of biological relevance with simple production protocols. However the methods for spheroid manipulation are still labor intensive, which severely limits the complexity of operations that can be performed on statistically relevant numbers of individual spheroids. Here we show how to apply hundreds of different conditions on spheroids in a single microfluidic chip, where each spheroid is produced and immobilized in an anchored droplet. By using asymmetric anchor shapes, a second drop can be merged with the spheroid-containing drop at a later time. This time-delayed merging uniquely enables two classes of applications that we demonstrate: (1) the initiation of cell-cell interactions on demand, either for building micro-tissues within the device or for observing antagonistic cell-cell interactions with applications in immuno-therapy or host-pathogen interactions, (2) a detailed dose-response curve obtained by exposing an array of hepatocyte-like spheroids to droplets containing a wide range of acetaminophen concentrations. The integrated microfluidic format allows time-resolved measurements of the response of hundreds of spheroids with a single-cell resolution. The data shows an internally regulated evolution of each spheroid, in addition to a heterogeneity of the responses to the drug that the single-cell analysis correlates with the initial presence and location of dead cells within each spheroid.

---

Recent years have seen the emergence of many new cell culture approaches to improve the relevance of *ex vivo* experiments to the behavior of the cells residing within living tissues. One of the main objectives of these methods is to recapitulate the native cells’ microenvironment, including biochemical signaling delivered from blood stream or from neighboring cells, the formation of intercellular junctions, interactions with the endogenous extra cellular matrix (ECM), mechano-transduction, and other effects such as diffusion gradients (1). The three-dimensional (3D) culture formats that have emerged range from the culture of individual cells in hydrogel matrices (2) or de-cellularized scaffolds (3), making functional aggregates such as spheroids (4) or organoids (5), or building more complex engineered structures that involve multiple cell types on a microfluidic device (6). Indeed the combination of microfluidics and 3D cell culture has allowed the emergence of a wide range of “organ on a chip” approaches that include many of these different strategies (7).

These formats are not meant to replace two-dimensional (2D) culture. Instead they will allow specific questions to be posed on more physiologically relevant culture models. Some of these questions can only be asked in specific 3D formats, such as questions related to embryogenesis (8), tumor-stromal interactions (9) or the effect of vascularization on tumor growth (10). In contrast, other applications depend on cellular phenotypes that are modified when the cells are cultured in 2D vs. 3D, such as the function of hepatocytes (11), chondrocytes (12), pancreatic (13), neural (14) or lung cells (15) and the impact of this function on their response to toxic compounds (16). Therefore the most suitable technological format for a particular question will balance the level of biological complexity that is required with the desired throughput and the necessary ease of use and reproducibility of the experiment.

In this context, spheroids present an appealing format for 3D culture, since they combine a moderately high level of biological complexity with simple production protocols (17). Indeed the biological function is enhanced in spheroids compared with 2D cultures (4), while cells have been shown to produce their own ECM and interact with it (18). However, despite the long history of spheroid cultures (19) and the ability to produce them in large quantities in bulk formats (20), the manipulation and observation of individual spheroids remains largely manual and labor intensive. As a result it is prohibitively difficult to link the bulk response, which is measured on the level of a population of spheroids, with the behavior of the individual cells within the spheroids and thus the 3D format itself.

In order to address these limitations we have demonstrated recently a microfluidic platform that integrates many of the necessary operations for the regulation of spheroids behavior *in vitro*, while providing several hundred independent cultures per experiment on a single microscope-slide format (21). The approach is based on using so-called “anchored droplets” (22) in which spheroids are formed, manipulated and observed over several days in culture. The ability to perform precise image analysis on the single-cell level, while combining results on thousands of spheroids, enables the mapping of cellular function depending on position within the spheroids, thus providing a link between the spheroid structure and the biological function of cells within it. The platform however could not address each spheroid individually with a specific condition and did not allow a succession of operations on them.

In the present paper we build on our previous results in order to allow random and time-dependent operations on each of the spheroids individually. This is achieved by introducing a new asymmetric design for the anchors, which leads to a qualitative transformation in the functionality of the microfluidic approach for a wide range of applications. Below, the physical principles of the devices and the protocols that allow combinatorial operations are first explained, followed by the description of two key classes of applications. First we describe the ability to bring cells into contact, for building complex tissues or to study antagonistic interactions between different cell types. These results have immediate applications in several areas of biological research such as tissue engineering, as models of immuno-therapies, or to understand host-pathogen interactions. Then we describe how the platform can be used to obtain a detailed drug dose-response on hepatocyte-like spheroids. The behavior is tracked by combining measurements of the time evolution of hundreds of spheroids with a single-cell resolution. The dynamics that emerges is fundamentally linked with the 3D structure of the spheroids and shows a strong effect of the cell-cell interactions on their response to the drug.

## Results

### Physical Principles of Differential Anchoring

In order to understand the principles underlying the device operation and robustness, we recall that confined droplets are subjected to a trapping force in regions where they reduce their surface area and thus their surface energy (22, 23). Therefore by designing microfluidic devices where the drops are confined everywhere, except in localized regions, as sketched in Fig. 1A, one can define positions at which the droplets can be anchored. The efficiency of this immobilization depends on the relative strength of the anchoring force, which is given by the gradient of surface energy, and the drag force due to the flow of the outer fluid: as long as the anchoring force is stronger than the drag force the droplet will remain immobile even if the outer fluid is flowing (23, 24).

**Fig. 1.**
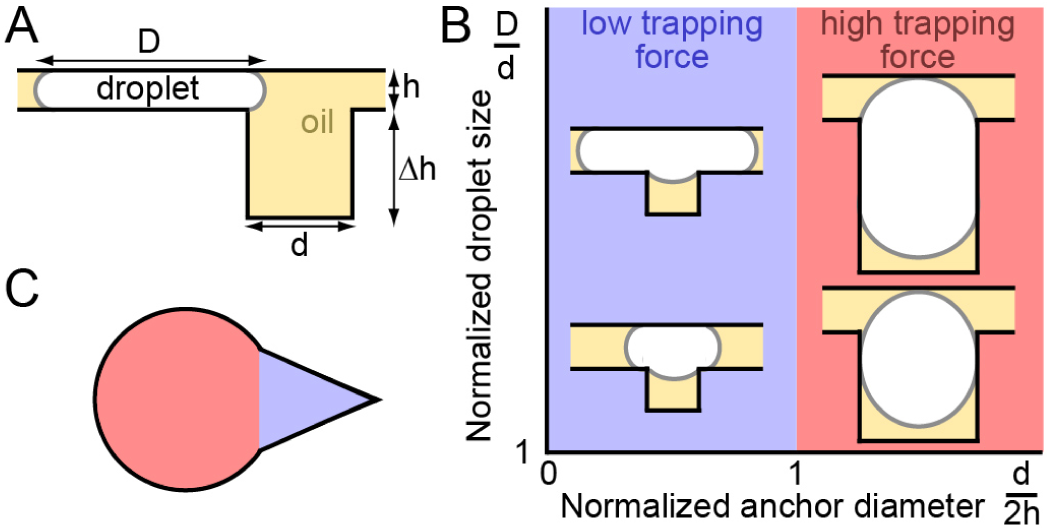
Physical principles of differential anchor strengths. (*A*) Side-view of a confined droplet near a capillary anchor. (*B*) Two anchoring strengths can be distinguished: for narrow anchors (blue shaded regions) the droplet enters only partially into the anchor, while for wide anchors (red shaded regions) the droplet enters entirely into the anchors. This leads to an anchoring efficiency that depends on the droplet size for the narrow anchors and to nearly irreversible trapping in the wide anchors. (*C*) The wide and narrow regions can be combined together into a single capillary anchor by designing asymmetric shapes.

When the droplet is above the anchor the curvature along the interface will tend to homogenize in order to equilibrate the Laplace pressure jump between the inside and outside. Therefore geometric considerations define two different limits that lead to two different regimes (Fig. 1*B*). First in the case of wide anchors, i.e. when the anchor diameter *d* is larger than 2*h*, the droplet penetrates completely into the anchor as long as the hole is sufficiently deep (23). This leads to a high trapping efficiency, as the large reduction of the surface area is combined with a weak drag force, since the droplet exposes only a small region in the channel where the fluid is flowing. Conversely, the droplet enters only partially into the anchor when *d* < 2*h*, leading to a critical flow velocity beyond which the anchor is not able to trap the droplet (22). Since larger droplets expose a larger cross-sectional area to the flow in this case, as sketched in Fig. 1B, the value of this critical velocity depends on the droplet volume, in addition to the physical parameters of the fluids (viscosity, surface tension). A detailed analysis shows that this critical velocity rapidly decreases as the droplet size increases (23) (see *SI Materials and Methods* for detailed discussion).

These principles can guide the design of anchors that have regions with different trapping efficiencies, as shown for example in Fig. 1*C*. The red-shaded region of this anchor displays a large diameter and so can accommodate a large droplet and trap it with a high efficiency, while still keeping the blue region free to receive a second droplet. In contrast, the largest dimension of this second region being smaller than the 2*h*, a second droplet will only be able to partially enter into the blue region, therefore being trapped with a weaker force. Depending on the design details of this shape the contrast between the two trapping efficiencies can indeed be very large, leading to nearly irreversible trapping in the red regions and much weaker trapping in the blue regions.

### Protocol for Droplet Pairing and Fusion

This differential trapping system can now be exploited to generate pairs of droplet having different contents. The protocol generally begins by bringing a population of large droplets and allowing them to randomly occupy the strong regions of the anchors, as shown in Fig. 2A (left). Once all of the anchors are filled, a second population of smaller droplets is transported into the trapping region, where they are trapped in the triangular parts of the anchors (right). With a slightly different anchor design, this protocol can be used to trap droplet multiplets Fig. S1. Beyond simply trapping the smaller drops, the triangular shape of the anchors also produces a local gradient of confinement that pushes the two drops in each anchor into intimate contact (27). As such, flushing an emulsion destabilization agent in the outer phase results in the quick merging of the two different types of droplets (Fig. 2*B* and Movie S1).

**Fig. 2.**
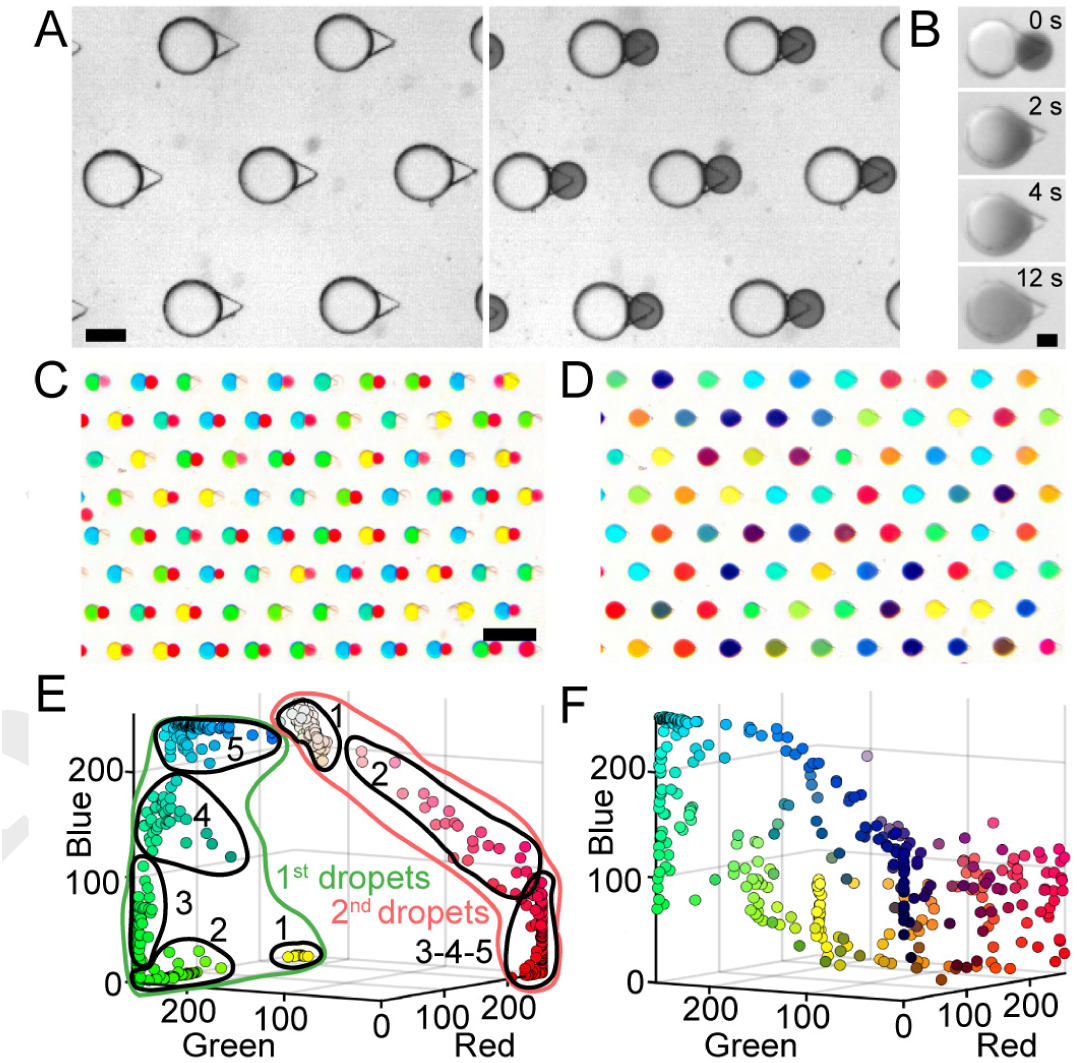
Protocol for pairing and merging different droplet populations. (*A*) First a population of large drops is injected and allowed to fill the large regions of each anchor, followed by a second population of smaller droplets. The small drops, which are colored with a dark dye, then occupy the triangular regions of each anchor. Scale bar is 200 *µ*m. (*B*) Flushing the device with an emulsion destabilization agent results in the merging of the touching droplets, which allows their contents to mix in a few seconds. Scale bar is 100 *µ*m. (*C-D*) Droplet libraries can be produced in a different microfluidic device and re-injected into the trapping region. In the current example, the large droplet population contains variable concentrations of dye ranging from blue to yellow, through different shades of green. The small droplets contain a gradient of red color. Image of 80 anchors filled with 2 sets of colored droplets before (*C*) and after (*D*) merging. Scale bar is 1 mm. (*E-F*) Quantification of the droplet colors in RGB space before (*E*) and after (*F*) the coalescence. The color of each dot in the 3D plot is given by its RGB coordinates (n_chip_ = 1; n_droplets_ = 351). Here the aqueous droplets are produced in fluorinated oil on the same microfluidic device that contains the anchoring region, using a modified flow-focusing junction (21) (see Fig. S2 for a typical chip design). All corresponding flows can be found in Table S1. The emulsion destabilization agent is PFO (e.g. perfluoro-octanol (25, 26)).

Alternatively one or both of the populations can be produced in a different device, stored off-chip, before being reinjected into the trapping region. In this way droplet libraries can be generated independently and later brought into contact with a sample of interest that is immobilized in the capillary anchors. Such a protocol is demonstrated in Fig. 2C-D, where two droplet populations, containing food dye as a proxy for chemical content, are merged together. In this example, each of the libraries was produced on a separate chip through a confinement gradient (28), as described in Fig. S3-4. The large drops were formed by mixing yellow and blue solutions, while the small drops contained a gradient of red dye. The large and small droplets were then sequentially loaded into the anchors (Fig. S5), to yield over 350 merged droplets each containing a unique color (Fig. 2E-F).

The demonstrations of Fig. 2 show the ability to bring together two droplets in each of microfluidic anchors. Below we show how this technology can be applied to cell manipulation, thus enabling unique operations on 3D cultures towards tissue engineering or screening applications.

### Constructive or antagonistic cellular interactions in anchored droplets

When a suspension of cells is encapsulated in the droplets, the aggregate together to form a single highly functional sphere of adherent cells in each anchored droplet (21). The size of the spheroid thus formed is determined by the number of cells in the droplet (Fig. S6). This protocol can be implemented in the current asymmetric anchors (Fig. 3*A*), resulting in a spheroid in the large droplet of each anchor. The second droplet can then be used to bring a different cell population that can interact with the original spheroid in a variety of ways, as illustrated in Fig. 3. Below a few examples of constructive or antagonistic interactions are described.

**Fig. 3.**
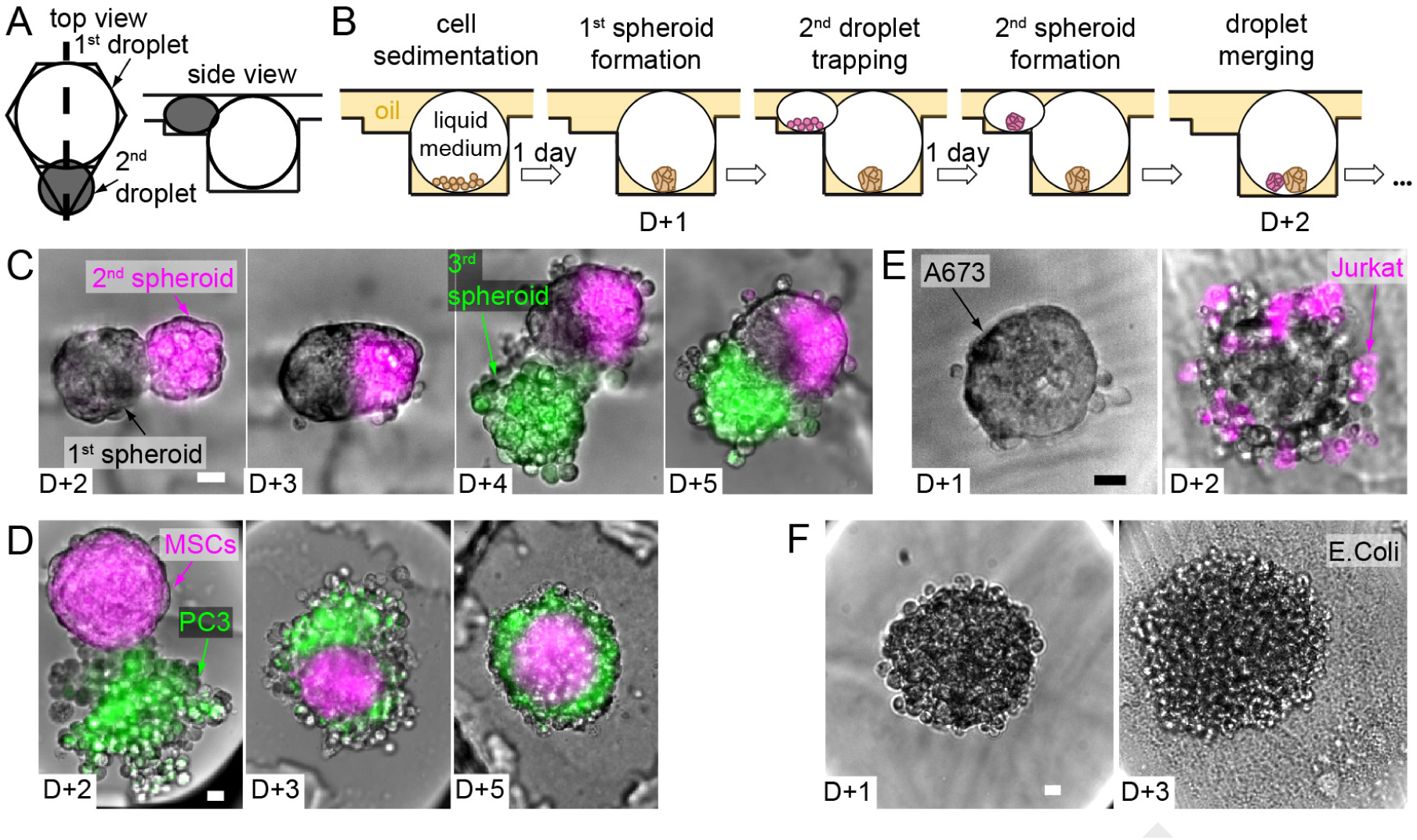
Spheroid merging and co-culture. (*A*) Design of the asymmetric anchors adapted for the droplet spheroid formation and culture (side view corresponds to the thick dashed line in the top view). (*B*) Scheme of the complete protocol for one spheroid merging. (*C-F*) Selected micrographs showing 2 consecutive merging events with H4-II-EC3 cell spheroids (*C*), and co-culture experiments with hMSCs - PC3 cells (*D*), Jurkat cells - A673 spheroids (*E*), and H4-II-EC3 cell spheroids - E.Coli (*F*). Scale bars are 20 *µ*m.

First, complex tissues can be constructed by successively bringing into contact cell populations. In the first example (Fig. 3*B-C*), a single H4-II-EC3 cell spheroid (a rat hepatoma cell line) was formed in each droplet trapped in the strong region of an anchor. The operation was repeated with a smaller droplet in the triangular region to form a single magenta-stained spheroid. After droplet pair merging, the two spheroids of each anchor came into contact in the resulting merged droplet and initiated fusion to form a composite spheroid (D+3). As the volume of the small droplet was much smaller than the large drop, the triangular region of the anchor was left empty after this first droplet merging. In this way, another trapping/merging cycle was possible, for instance with a new set of droplets containing cells stained in green (Fig. S7*A*). This sequential three step process resulted in the formation of a single composite microtissue in each anchor of the microfluidic chamber. Since all of the spheroids were formed by the same cell type, these composite spheroids displayed a random organization of the differently stained cells in each anchor, resulting in a wide variety of final shapes (Fig. S7*B*).

In contrast, the microtissues organized into well organized structures when they involved different cell types, as shown in Fig. 3*D*. Here the large droplet contained a spheroid of human mesenchymal stem cells (hMSCs) and the smaller droplet contained an aggregate of PC3 cells (a prostate cancer model), as a tumor-stroma interaction model (29). In this case the hetero-spheroid always took a core-shell structure, with the hMSCs occupying the core of the spheroid, while the PC3 cells formed a shell. Such a core-shell model was reproduced in each of the anchors, independent of the details of the initial cell configurations.

In addition to these constructive interactions, the droplet merging can also be used to explore host-pathogen or other antagonistic interactions between different cell types. For instance, the microfluidic approach is well suited for studying the interaction of immune cells with a cancer spheroid *in vitro*. A simple example of such an interaction is obtained by bringing into contact a spheroid of A-673 cells (Ewing’s sarcoma) with a suspension of individualized Jurkat cells (Fig. 3*E*). These immune cells form robust junctions with the cancer spheroid and begin to interact with the cancerous cells in each of the anchors.

Finally, other host-pathogen interactions can be explored by encapsulating bacteria in the second droplet and bringing the bacteria into contact with the initial spheroid. This is exemplified in Fig. 3*F*, where a well-formed spheroid (H4-II- EC3 cell) is brought into contact with a droplet containing a dilute suspension of *E. Coli*. Two days after the merging, the bacterial colony has exploded and the toxic effect of the bacteria on the mammalian cells is apparent through the dismantling of the spheroid.

The examples above are only meant to illustrate the range of interactions that are possible in the device. These operations can be generalized to many other cell types and interactions. For instance the timing of droplet merging can be varied with respect to the timing of spheroid formation, providing a way to investigate the self-assembly of a well-formed spheroid with a non-reorganized aggregate (Fig. S8). Moreover the introduction of a hydrogel into the one or both of the droplets can allow cells to interact together through paracrine signaling while remaining physically separated by a porous barrier in same the droplet (Fig. S8).

### Dynamic Measurement of Drug Toxicity on Spheroids

Beyond cell-cell interactions, the microfluidic platform also lends itself to screening different chemical conditions in each droplet. As a demonstration of the type of approach that can be applied, we show how the asymmetric anchors can be applied for measuring the concentration-dependent acetaminophen toxicity (APAP, a drug known for its hepatotoxicity (30)) on H4-II-EC3 cell spheroids (Fig. 4*A-B*). For this purpose, the spheroids were formed in the large droplets, as shown above (Fig. 4*C*). In parallel a library of small droplets containing a range of APAP concentrations was prepared as described in *SI Materials and Methods*. The APAP stock solution was marked with a magenta fluorescent probe, while the dilutant solution contained a green fluorescent probe. In this way, the two fluorescent signals could be used to determine the APAP concentration in each of the droplets (Fig. S9). Droplets without drugs or label were also produced in order to obtain control conditions in some of the anchors. Then these drug droplets were injected into the chip, trapped alongside the spheroids (Fig. 4*D*), and merged with them. Consequently, the spheroids on a single device were exposed to range of APAP concentrations covering three decades (Fig. 4*E*) and the evolution of cell death was dynamically monitored for 36 h in each spheroid by live viability staining.

**Fig. 4.**
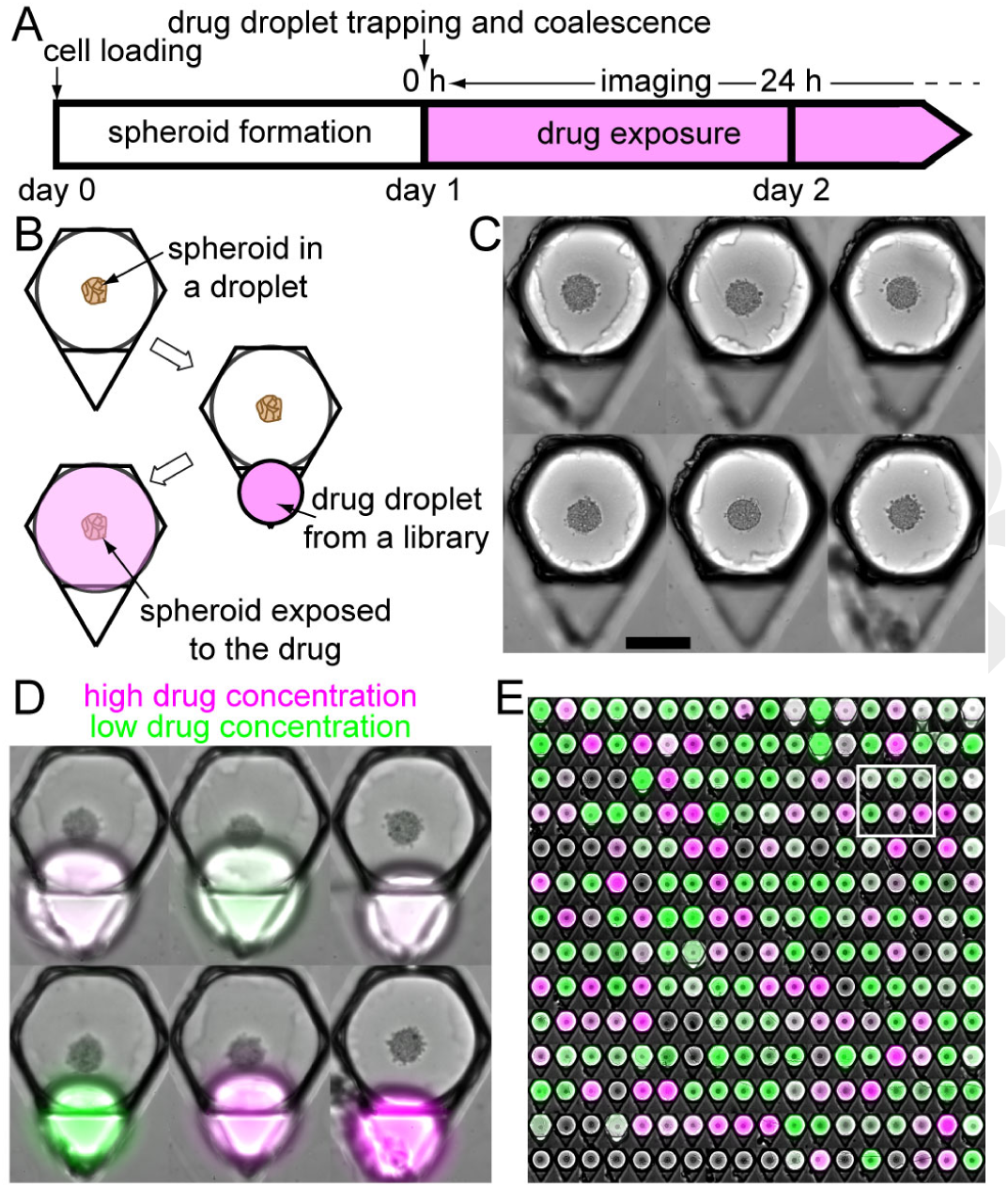
Drug toxicity experiment on liver spheroids. (*A-B*) Timeline (*A*) and schematic top view of an anchor (*B*) showing the experimental protocol. (*C-E*) Montages of 6 micrographs showing anchors with single liver spheroids (H4-II-EC3 cells) before (*C*, scale bar is 200 *µ*m) and after (*D*) the drug droplet trapping, and of the entire chip array after droplet coalescence (*E*, n_spheroids_ = 252). The green and magenta fluorescent dyes corresponds respectively to the APAP stock solutions at low and high concentrations, used for creating the droplet library. The white rectangle in (*E*) shows the location of the 6 anchors displayed in (*C-D*).

The results at the low and high extremes of the concentration range were as expected: spheroids that were exposed to control droplets or low APAP concentrations remained viable for the whole duration of the experiment (Fig. S10), while the viability and cohesiveness of the spheroids exposed to high APAP concentrations was altered (Fig. 5*A*). Indeed, the mean viability at 24 hours displayed the typical sigmoidal shape on a logarithmic scale (Fig. 5*B* and Fig. S11) with a half maximal inhibitory concentration (IC50) of 18.0 mM (a similar value is found in 2D, Fig. S12). The variability between different microfluidic chips was very low since the IC50 coefficient of variation was only 3 % (n_chips_ = 4).

**Fig. 5.**
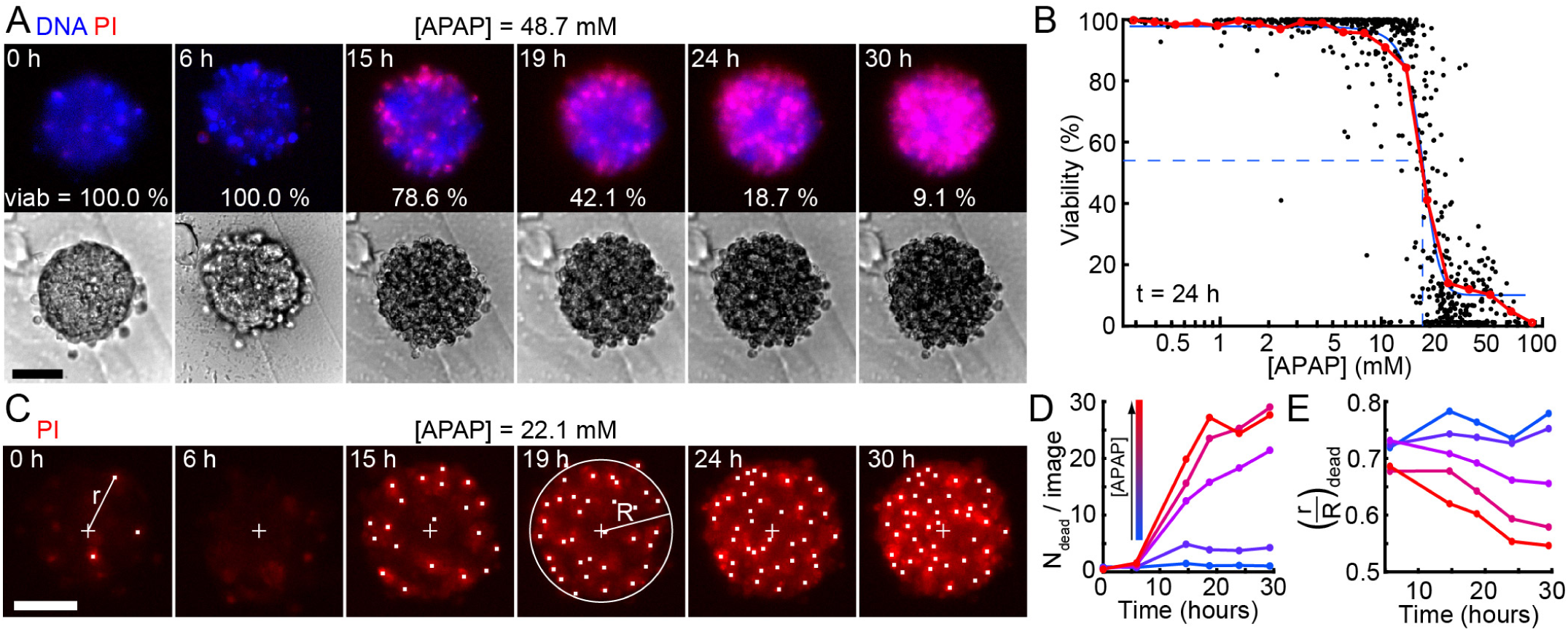
Acetaminophen (APAP) toxicity on H4-II-EC3 cell spheroids. (*A*) Time lapse images showing a spheroid exposed to a 48.7 mM APAP concentration, in bright field (bottom) and with the fluorescent viability staining (top). (*B*) Toxicity values at the spheroid level after a 24 h exposure (n_spheriods_ = 685). Each black dot represent one spheroid, the red and blue curves represent respectively the mean behavior and a sigmoidal fit of the data, with the blue dashed lines highlighting the IC50 value of 18.0 mM. (*C*) Time lapse images showing a spheroid exposed to a 22.1 mM APAP concentration with the mortality marker (propidium iodide, red). White dots are the locations of the detected dead nuclei and the cross represents the spheroid center. R is the equivalent radius of the spheroid and r is the distance to the spheroid center. (*D-E*) Time evolution of the number of dead cells detected on one spheroid image (*D*) and of the mean normalized distance 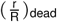 of the dead cells to the spheroid center (*E*) depending on the drug concentration. Blue to red: [APAP] < 5 mM, n_spheriods_ = 203; 5 mM < [APAP] < 15 mM, n_spheriods_ = 215; 15 mM < [APAP] < 23 mM, n_spheriods_ = 98; 23 mM < [APAP] < 40 mM, n_spheriods_ = 127; 40 mM < [APAP], n_spheriods_ = 53. Scale bars are 50 *µ*m.

More interestingly, the experiments yielded a deeper understanding of the drug response when the time evolution of each spheroid was tracked at the level of individual cells (Fig. 5*C*). For this purpose, we detected the apparition of dead cells as function of time (Fig. 5*D*) and measured their distance to the spheroid center for each of the 685 spheroids 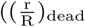, see *SI Materials and Methods*, Fig. 5*E*, individual curves shown in Fig. S13). At low APAP concentrations, the number of dead cells remained below five cells per image and their mean location in the spheroid was constant and close to the spheroid edge 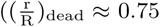, light and dark blue lines). For higher APAP concentrations, the number of dead cells increased significantly after 10 h to reach up to 30 dead cells at the end of the experiment, corresponding to all the cells on an epifluorescence image. In addition, the position of these dead cells shifted towards the spheroid center with time and concentration, with 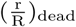 dropping from 0.68 to 0.55 for an APAP concentration higher than 40 mM (red line). This indicated that the drug concentration influenced the number and also the location of dead cells within the spheroids, in a time dependent manner.

The single-cell level of detail allowed us to address the wide spread that was observed for intermediate concentrations of APAP. Indeed, for APAP concentrations between 15 and 23 mM, spheroids could have a very low, very high or intermediate viability (Fig. 6*A*). One major parameter for explaining this spread was the presence of dead cells in the spheroids at the outset of the experiment. Indeed, spheroids with at least one dead cell at t = 0 h displayed, at t = 24 h, a significantly lower viability (24 %) than the spheroids without initial dead cells (60 %, Fig. 6*B*). Moreover, the location of these first detected dead cells was significantly correlated to the viability after 24 h (Fig. 6*C*): the spheroid was more likely to have a low viability at t = 24 h when the first dead cell was close to the spheroid center 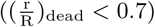.

**Fig. 6.**
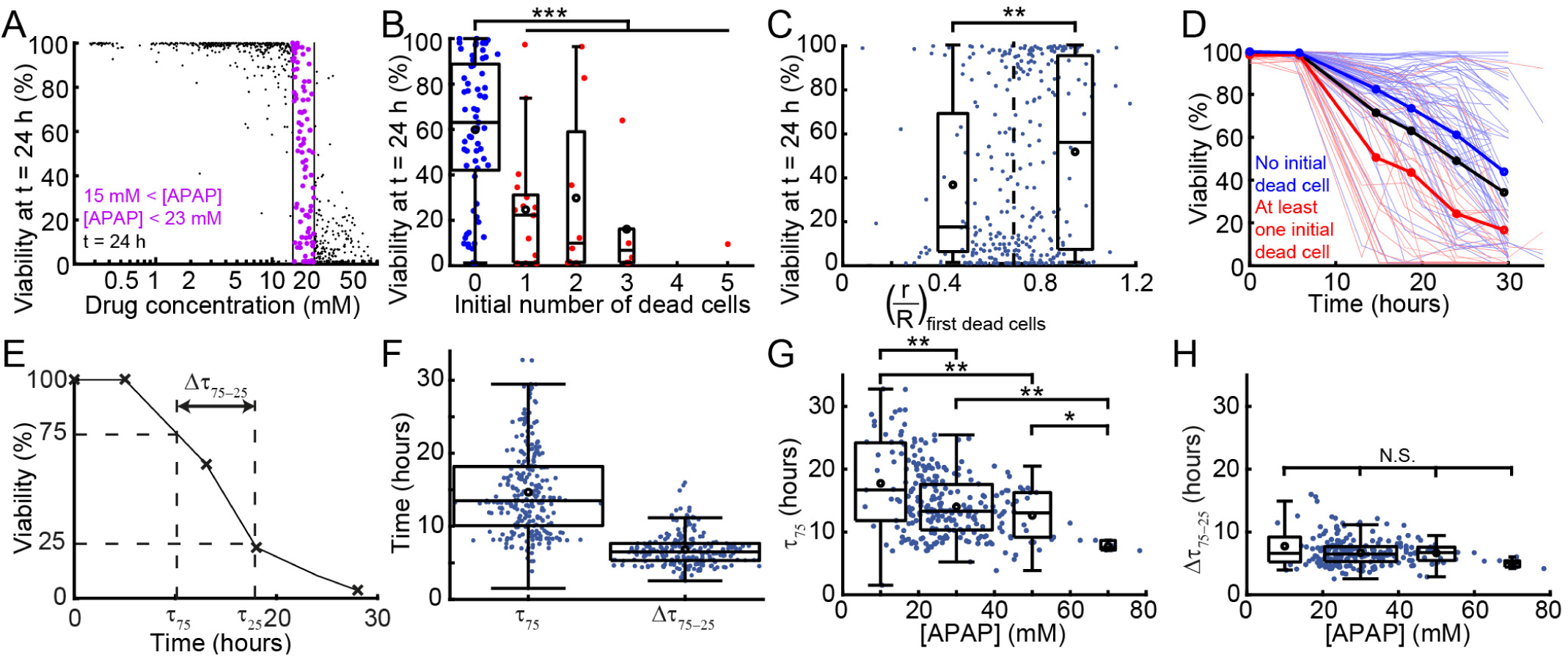
Analysis of the viability dispersity. (*A*) Toxicity values after a 24 h exposure with the data points corresponding to spheroids exposed to an APAP concentration between 15 and 23 mM highlighted in purple. (*B*) Influence of the number of initial dead cells on the viability after a 24 h exposure, for an APAP concentration between 15 and 23 mM (red: at least one initial dead cell, blue: no initial dead cell, n_spheriods_ = 98). (*C*) Correlation between the viability at t = 24 h and the mean normalized distance of the first detected dead cells (whatever its time of appearance) to the spheroid center 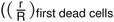, n_spheriods_ = 308), for an APAP concentration between 10 and 40 mM. (*D*) Dynamic evolution of the spheroid viability for an APAP concentration between 15 and 23 mM. Each thin line represents one spheroid (n_spheriods_ = 98), the red and blue curves correspond respectively to the spheroids that had at least one detected dead cell and no detected dead cell at t = 0 h while the thick black line represents the overall mean. (*E*) Definition of the time needed to reach a 75 % viability (*τ*_75_) and the time to go from a 75 % to a 25 % viability (Δ*τ*_75-25_ = *τ*_25_ – *τ*_75_) on a viability followup corresponding to a spheroid exposed to a high APAP concentration (above 40 mM). (*F*) Evaluation of *τ*_75_ (n_spheriods_ = 262) and Δ*τ*_75-25_ (n_spheriods_ = 215). (*G-H*) Evolution of *τ*_75_ (*G*) and Δ*τ*_75-25_ (*H*) with the APAP concentration. N.S.: non significant; *: p < 0.05; **: p < 0.01; ***: p < 0.001.

The signature of the initial state was again visible when observing the time-evolution of the viability of each spheroid (Fig. 6*D* and Fig. S14), where the curves displayed different trends depending on the presence or absence of a dead cell initially. This observation motivated us to question what part of the viability dynamics was determined by the drug concentration and what part depended on the *eigenstate* (the structure, interactions and initial spheroid state) of each spheroid in time. In order to evaluate the different effects we defined two time scales to describe the evolution of each spheroid, as sketched in Fig. 6*E*: the time to reach 75 % viability (*τ*_75_) and the time required to drop from 75 % to 25 % viability (Δ*τ*_75-25_). The first time scale could be considered as the time required for the cells to begin responding to the drug, while the second one described the time required to kill the spheroid once the process began.

These two parameters were found to be independent of each other (Fig. S15). They also had different mean values and variances (Fig. 6*F*), such that Δ*τ*_75-25_ could be considered as nearly constant compared to *τ*_75_. Indeed, *τ*_75_ showed a significant dependence on APAP concentration, dropping from 18 to 8 h for APAP concentration from 10 to 70 mM (Fig. 6*G*), while Δ*τ*_75-25_ remained constant close to 7 h for all concentrations (Fig. 6*H*). These measurements suggested that *τ*_75_ was the more relevant parameter to understand the effect of the drug, while Δ*τ*_75-25_ was characteristic of the response of these spheroids to a change in their microenvironment.

## Discussion

As the demand for highly relevant culture models becomes stronger (31), many approaches have been explored for structuring 3D cultures that capture essential aspects of the *in vivo* micro-environment (32). Among these approaches, spheroids constitute an interesting format that balances a very good biological relevance while remaining simple to produce in large quantities. Nevertheless, spheroid manipulation remains laborious and technical, which severely limits the ability to generate large data sets on a variety of culture conditions (17). In this context we have recently demonstrated the ability to obtain multiscale cytometry by performing phenotypic measurements on 10^5^ – 10^6^ individual cells *in situ* within thousands of spheroids (21). Here we complete the toolbox by developing a protocol to manipulate independently each of the hundreds of spheroids present on a microfluidic device, which allows each spheroid to be subjected to a particular treatment upstream of the data collection.

These operations hinge on the ability to reliably pair two droplets together, at any moment over a period of several days. The microfluidic design uses anchors that contain regions with two widely varying trapping strengths, such that the primary droplet can be trapped nearly irreversibly while other droplets are brought into contact at later times. This contrast in trapping efficiencies makes the device operation possible for a wide range of physical and experimental parameters (viscosities, flow rates, droplet sizes, etc.), which translates into much stronger robustness and stability compared with existing designs (e.g. (26, 27, 33, 34)). These qualities bring droplet microfluidics closer to the functionalities of multiwell plates in which any multi-step chemical or biological assay is possible (35). When compared with the 96 well plate however, the volume in each of our droplets is about 1,000 times smaller and the number density of conditions per cm^2^ in our chip is 100 times higher. Moreover the integrated microfluidic format allows operations, such as droplet loading and merging, to be performed in parallel in all positions, which removes the need for bulky pipetting robots.

When applied to spheroid cultures, the resulting protocols provide a way to bring the contents of different droplets into contact while respecting the biological time scales of the cells. As such the platform allows the user to wait until after the spheroids have formed before adding new contents to the primary droplets. This new content can consist of cells of different types, for example to build micro-tissues within the droplets (36), to study cell-cell interactions (37), to reproduce crucial steps in development (38), or in order to study antagonistic interactions as in immuno-therapy models (39) or host-pathogen interactions (40). Although all of the above subjects are of strong current interest, they all lack a robust and high-throughput platform for performing the 3D cultures, relying instead on manual operations. In this context, the experiments performed in this study illustrate the types of interactions that can easily be explored with the current format. The large number of parallel operations, along with the integrated format, couple naturally with any microscopy platform to generate very large data sets with a single-cell resolution. In turn this provides a way to quantify the heterogeneity of cellular interactions and to search for rare events that may have a significant impact *in vivo*. For instance, future work with more relevant immuno-therapy models will allow us to quantify these interactions in hundreds of parallel droplets in order to characterize the heterogeneity of immune cell response. Alternatively, tracking the motion of each of the immune cells can indicate if the cells selectively target parts of the tumor spheroid.

Beyond these illustrative examples, the ability to manipulate individual spheroids allows us to study the toxicity of a drug (acetaminophen) on hepatocyte 3D cultures. A complete screen is reported here, including generating the droplet library, merging the library with the spheroids, following the response of individual cells within hundreds of spheroids, and analyzing the results at the single-cell level and as a function of time. The results obtained in this section are noteworthy on several levels:

– First as a demonstration of the strength of the technology, since such detailed data would have been prohibitively difficult to obtain using the current state of the art (41–43). In contrast, the integrated microfluidic format made the experiments simple to perform and the measurements straightforward.
– Second, the large number of spheroids involved in the study highlighted the presence of a strong heterogeneity of responses near the IC50 position, where some spheroids were completely viable and others were completely dead after 24h. By using the single-cell longitudinal measurements, this variety of responses was found to be correlated to the presence of dead cells within the spheroids at the initial moment, indicating that the structural integrity of the spheroid played a role in its ability to resist a drug treatment.
– Third, the viability measurements showed complex dynamics, with the emergence of two time scales from the time-resolved single-cell measurements: *τ*_75_ that depends on the drug concentration and Δ*τ*_75-25_ that does not. These observations are consistent with the idea that once a spheroid has reached a 75% viability, it will proceed to entirely die at a rate that is proper to the spheroid and independent of the external stimulus. Indeed if we consider a particular cell whose neighbors die due to the drug, leading it to loose its focal adhesions and to be subjected to toxic hydrolases, the change in its micro-environment can induce (or at least precipitate) its death for purely structural reasons. The presence of such a mechanism is supported by the bright field images, which show that the spheroids loose their cohesiveness as the cells within them die.

While the results discussed here do not address the question of whether spheroids are better predictors of in vivo drug response than other formats (44–46), they do indicate that the spheroid response to a drug involves collective structural effects that are not present in 2D monolayers. Such effects play a role in the response of solid tumors to chemotherapy and may provide important insights on the sensitivity or survival of micro-niches within a tumor, in particular as cancer cells may be interspersed with fibroblasts or other stromal cell types. The link that appears between the structure and biological response within a spheroid thus illustrates ways in which 3D cell culture captures biological complexities in ways that 2D cultures cannot.

Looking ahead, the technological tools presented here can also be transposed to other 3D culture formats including organoids (47), blastoids (48), or embyoid bodies (49), following some adaptations of the protocols. In addition to this, it is straightforward to combine the different operations, such as building tissues in the microfluidic anchors and then testing the effects of different molecules on these hetero-spheroids, or testing combination treatments involving small molecules, cellular therapy, and other approaches in a single device. Finally, the secondary droplet can also be used to induce a phase change from liquid to solid in the primary droplet, either by introducing a hydrogel (matrigel, agarose, etc.) or by introducing a cross-linking agent into a droplet that already contains the gel. Such hydrogels can then be used for a wide range of applications e.g. for metastasis invasion assays (9) or as functionalized scaffolds to capture secreted molecules (2).

Finally, as stated in the introduction, the most suitable 3D model depends on the question being addressed. So future work will involve a careful choice of the cell types and combinations, the 3D culture format, as well as a validation steps to ensure that the results obtained from the model system are relevant to the *in vivo* conditions. The microfluidic platform described here is designed to make such studies and validation much more efficient and reproducible.

## Materials and Methods

### Estimation of the trapping and drag forces

Let us consider the anchors described in the main text in Fig. 1 and Fig. 2. The microfluidic channel has a height *h* and the circular parts of these anchors have a diameter *d* and a depth Δ*h*. In order to estimate the trapping force of the first droplets in the circular parts of the anchors, the variation of surface area before and after trapping must be estimated. The surface area of a confined droplet is estimated by considering pancake shape, meaning a cylinder of radius *R*_*i*_, height *h* surrounded by the outer half of a torus of small radius 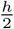. As we consider the case where the droplet size matches the circular part volume of the asymmetric anchor, the surface area of the trapped droplet is estimated by calculating the area of a cylinder whose section is the circular part of the anchor and whose height is *h* + Δ*h*. Thus, the trapping force of the first droplets in the circular parts of the anchors is:

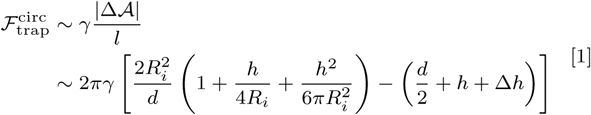

where the characteristic length *l* over which the surface energy changes is estimated by 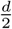.

For estimating the trapping force of the first droplets in the triangular parts, the triangular parts are assimilated to small circular anchors, whose radius 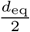 is smaller than the channel height. According to Dangla *et al.* (23), the resulting trapping force is:

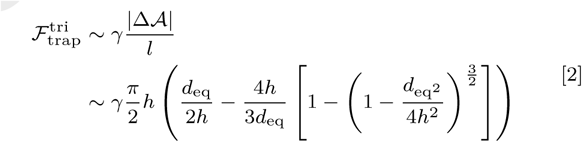

With the experimental parameters used for Fig. 2 (*R*_*i*_ = 125 *µ*m, *d* = 250 *µ*m, *h* = 95 *µ*m, Δ*h* = 50 *µ*m) and assuming *d*_eq_ = 150 *µ*m (equal to the length and width of the anchor triangle), we have:

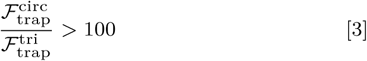

So, for the first droplets, the trapping in the circular parts of the anchors is much more efficient than the trapping in the triangular parts.

Also, the drag force exerted by the fluid on a confined droplet scales as 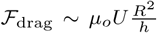, where *U* and *µ*_*o*_ are respectively the mean velocity and viscosity of the outer fluid. Therefore, the ratio between the drag force exerted on the large first (*R*_1_ = 170 *µ*m) and small second (*R*_2_ = 80 *µ*m) confined droplets before trapping is:

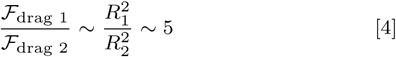

Consequently, if the small droplets experience a similar trapping force in the triangular parts of the anchors than the large droplets, they are exposed to a smaller drag at constant flow rate, meaning that their trapping in the triangular parts of the anchors is more robust.

### Microfabrication

Molds were mainly fabricated using standard dry film soft lithography. Up to five layers of dry film photoresist, consisting of 50 and 33 *µ*m Eternal Laminar (respectively E8020 and E8013, Eternal Materials, Taiwan) and 15 *µ*m Alpho NIT215 (Nichigo-Morton) negative films, were successively laminated using an office laminator (PEAK pro PS320) at a temperature of 100 °C until the desired channel height, from 50 to 200 *µ*m depending on the different cases, was reached. After each laminating step, the photoresist film was exposed to UV (LightningCure, Hamamatsu) through a photomask of the junction, channels, trapping chamber boundaries or anchors. The masters were revealed after washing in a 1 % (w/w) K_2_CO_3_ solution (Sigma-Aldrich). For the 3D anchors fabrication (Fig. 3 to Fig. 5), a specific method was developed. In these cases, the top of the chip consisted of the flow-focusing device and chambers and the anchors were located at the bottom of these chips. The anchors mold was designed with RhinoCAM software (MecSoft Corporation) and was fabricated by micro-milling a brass plate (CNCMini-Mill/GX, Minitech Machinery). That was also the case for the droplet library producing chips with an aqueous injector and a slope (see Fig. S3). The topography of the molds and masters were measured using an optical profilometer (VeecoWyco NT1100, Veeco). For the fabrication of the top of the chip, poly(dimethylsiloxane) (PDMS, SYLGARD 184, Dow Corning, 1 g of curing agent for 10 g of bulk material) was poured over the master and cured for 2 h at 70 °C. For the 3D anchors, the metallic mold was first covered with PDMS. Then, a glass slide was immersed into uncured PDMS, above the anchors. The mold was finally heated on a hot plate at 180 °C for 15 minutes before extraction of the glass slides covered by a thin layer of PDMS with the anchor pattern. In all cases, the top and the bottom of chip were sealed after plasma treatment (Harrick).

### Chip Design

Two main different chip designs were used in this study, depending on the presence of cells. For the experiences involving cells (Fig. 3 to Fig. 5), the chip design is detailed in Fig. S2. In this case, there were 252 anchors disposed along an hexagonal pattern in the 2 cm^2^ trapping chamber. For the non biological experiments (Fig. 2), the design was similar but with different dimensions. Notably, the heights before and after the step at the junction were respectively 50 *µ*m and 83 *µ*m. Rails allowed the droplets to be distributed homogeneously across the chamber, that had a 95 *µ*m height. In this case, contrary to the cellular chip, the 50 *µ*m deep anchors were patterned on the top of the chamber.

### Experimental Microfluidic Protocol

The chips were filled 3 times with Novec Surface Modifer (3M), a fluoropolymer coating agent, for 30 min at 110 °C on a hot plate. All experiments were conducted using the FC40 fluorinated oil (3M) implemented with a biocompatible FluoroSurfactant (Ran Biotechnologies) at different concentrations. The solutions were loaded in glass (SGE) or plastic (Terumo) syringes, that were actuated with programmable and computer controlled syringe pumps (neMESYS, Cetoni). The syringes were directly connected to the PDMS chips with PTFE tubing (Adtech). For the merging of droplet pairs, the trapping chambers were perfused with a 20 % (v/v) 1H,1H,2H,2H-perfluoro-1-octanol (Sigma-Aldrich) solution dissolved in Novec™-7500 Engineered Fluid (3M) at the flowrate indicated in Table S1. The uncolored and dark droplets seen in Fig. 2 and Fig. S1 are respectively made of pure water and of a 6 mM 2,6-dichlorophenolindophenol (2,6-DCPIP, Sigma-Aldrich) aqueous solution.

### Cell and Bacteria Culture

A rat H4-II-EC3 hepatoma cell line (American Type Culture Collection, CRL-1600™, LGC) and A-673, an muscle Ewing’s Sarcoma cell line (ATCC, CRL-1598™, LGC) were maintained on T-25 cm^2^ flasks (Corning) in a standard CO_2_ incubator (5 % (v/v) CO_2_, C150 incubator, Binder), following the instructions provided by the manufacturer. The culture medium was composed of Dulbecco’s Modified Eagle’s Medium (DMEM, ThermoFischer) containing high glucose supplemented with 10 % (v/v) fetal bovine serum (ThermoFischer) and 1 % (v/v) penicilin-streptamicine (ThermoFischer). The cells were seeded at 5.10^4^ cells/cm^-2^ and sub-cultivated every 3 days.

Human mesenchymal stem cells derived from the Wharton’s jelly of umbilical cord (UC-hMSCs) (ATCC, PCS-500-010, lot #63516504, LGC) were obtained at passage 2. UC-hMSCs were maintained in T-175 cm^2^ flasks (Corning) and cultivated in a CO_2_ incubator. The culture medium was composed of Alpha Modified Eagle’s medium (*α*-MEM) (Gibco, ThermoFischer) supplemented with 10 % (v/v) fetal bovine serum (Gibco) and 1 % (v/v) penicilin-streptamicine (Gibco). The cells were seeded at 5.10^3^ cells/cm^2^, sub-cultivated every week, and the medium was refreshed every 2 days. UC-hMSCs at passage 2 were first expanded until passage 4 (for about 5-6 populations doublings, PDs), then cryopreserved in 90 % (v/v) FBS / 10 (v/v) % DMSO and stored in a liquid nitrogen tank. The experiments were carried out with UC-hMSCs at passage 4 to 8 (about 24-35 PDs, after passage 2).

PC-3, a human prostate adenocarcinoma cell line (ATCC, CRL- 1435™, LGC) was maintained on T-75 cm^2^ flasks (Corning), and Jurkat, a human lymphoblast cell line (ATCC, TIB-152™, LGC) was was maintained in suspension using T-25 cm^2^ flasks (Corning) in a standard CO_2_ incubator (5 % (v/v) CO_2_), following the instructions provided by the manufacturer. For both cell types, the culture medium was composed of Roswell Park Memorial Institute medium (RPMI, ThermoFischer), supplemented with 10 % (v/v) fetal bovine serum (ThermoFischer) and 1 % (v/v) penicilin-streptamicine (ThermoFischer). The PC-3 cells were seeded at 5.10^4^ cells/cm^2^ and Jurkat were seeded at 1.10^5^ cells/mL. Both cell lines were sub-cultivated every 3 days.

*Esherichia Coli* (K12 - MG 1655 strain) were cultivated as colonies on LB-agar plates. The day of the fusion experiments, *E. Coli* were cultivated in suspension using LB medium up to reaching an O.D. of 0.5.

### Spheroid Formation on Chip

The chips were first filled with a 3 % (w/w) FluoroSurfactant solution. All air bubbles were discarded. H4-II-EC3 cells were detached from the culture flasks with a 2-3 minutes incubation in TrypLE™ Express enzyme (ThermoFischer), that was then inactivated by addition of warm medium. The resulting cell solution was centrifuged (centrifuge 5702 R, Eppendorf) at 2,400 rpm for 6 minutes while the cell concentration was determined using a haemocytometer (Marienfeld). The supernatant was discarded and the cell pellet was resuspended at a 6.10^6^ cells/mL for direct use, or 8.6.10^6^ cells/mL before addition of agarose, in culture medium supplemented with gentamicin (Sigma-Aldrich) to a final concentration of 50 mg/L. When needed, the agarose stock solution was prepared in parallel. Ultra-low gelling agarose (Type IX-A, Sigma-Aldrich) was dissolved at a 3 % (w/w) concentration in warm sterile PBS implemented with gentamicin to a final concentration of 50 mg/L and kept at 37 °C. 30 *µ*L of the agarose stock solution and 70 *µ*L of the cell solution were mixed to obtain a final cell concentration of 6.10^6^ cells/mL in a 0.9 % (w/w) agarose solution. One glass syringe was loaded with this solution and droplets were produced according to the flowrates displayed in Table S1. Spheroids of hMSCs, A-673, PC-3 and H4-II-EC3 were formed in droplets containing DMEM medium, while Jurkat were loaded on chip in RMPI medium. The cell loading was performed at 37 °C in a microscope incubator (Okolab) in which all chips, syringes, connectics and solutions were pre-heated. After the loading, all flowrates were stopped, the tubings were cut and the chips were kept immersed in PBS in the CO_2_ incubator. Cells started sedimenting at the bottom of each droplet when the flowrates were stopped. They reorganized overnight in the liquid agarose droplets into spheroids. For the toxicity experiments (Fig. 4 to Fig. 5), the gelation allowed to immobilize the spheroids at the bottom of their droplets, facilitating live imaging.

### Spheroid Preparation and Staining for Co-culture Experiments

A 10 mM solution of CellTracker™ Red and Green (ThermoFischer) was prepared in sterile DMSO (PAN Biotech). H4-II-EC3 cells were incubated for 30 min in culture medium with 10 *µ*M of CellTracker™ in the culture flask, before PBS washing and exposition to the TrypLE™ express enzyme. Alternatively, Jurkat and hMSCs were stained for Vybrant™ Dil (red) and PC-3 were labeled with Vybrant™ Dio (greeen), following the manufacturer instructions (ThermoFisher).

### Protocol For Droplet Library Production

Droplet libraries were produced following two successive steps. First, the two solutions of interest were mixed at a PEEK cross junction (Upchurch) in known ratios controlled by programmable syringe pumps (neMESYS, Cetoni). Between each different ratio, a plug of oil (FC40 with a 0.25 % (w/w) concentration of FluoroSurfactant) was injected at the cross junction for physically separating the different droplets in the exit tubing. This technique is called micro-segmented flow (50) and results in a train of microliter droplets separated by oil, each of them with a different ratio of the two aqueous stock solutions (see Fig. S4*A-B*). In the present study, the ratios followed either a linear (Fig. 2, Fig. S4-5) or logarithmic (Fig. 4 and Fig. 5) progression.

Second, these segments were partitioned into nanoliter droplets using a specific droplet producing chip (Fig. S3). The segments were injected in a chamber filled with oil through a slope (28) (see Fig. S3*A*). The slope allows to continuously deconfine the aqueous phase that spontaneously breaks into monodisperse droplets, without the need of an external oil flow (Fig. S3*B*). The droplet size is governed by the geometrical parameters of the injector, namely its height and width, as well as the angle of the slope. The nanoliter droplets resulting from this production were brought in the storage chamber of the chip thanks to a small continuous oil flow (FC40 with 6 % (w/w) FluoroSurfactant) at the corner of the slope (Fig. S3*C-D*). The droplets ascended to the top of this very deep storage chamber resulting in the trapping of the produced droplet library.

### Food Dye Droplet Library Production

The color droplets shown in Fig. 2, Fig. S4 and Fig. S5 were produced by mixing commercially available yellow, blue, red food dyes (Vahiné) and pure water in known ratios. The syringe pumps were programmed to create 11 segments (5 with a linear increase of the first solution, 1 purely made of the first solution, and 5 with a linear decrease of first solution), each of 2 *µ*L, separated by 1 *µ*L of oil at a global constant flowrate of 20 *µ*L/min. To partition these segments at the desired volumes, two droplet producing chips were used. For the first and second droplets, the geometrical parameters of the chip were respectively a slope of 8 and 11 %, an injector width of 100 and 90 *µ*m and an injector height of 40 *µ*m for both. The flowrates are indicated in Table S1.

### Acetaminophen Droplet Library Production

Two stock solutions were prepared for the acetaminophen (APAP, Sigma-Aldrich) droplet library production, with a 3 mM and 300 mM drug concentration. Both of them were implemented with DMSO (8 % (v/v)) and gentamicin (50 mg/L). For the live viability staining, the 2 solutions of the manufacturer (ReadyProbes™ Cell Viability Imaging Kit (Blue/Red), ThermoFischer) were diluted to a 17.5 % (v/v) concentration. The low and high APAP drug concentration solutions were marked respectively with a green and red fluorescent dye (CF™ 488A hydrazide and CF™ 647 hydrazide, Sigma-Aldrich) at a 7 *µ*M concentration. The dilutant was culture medium. These fluorescent dyes did not have any observable effect on the spheroids viability at the highest concentration used in this study. A solution was also prepared for the control droplets with the same composition as the drug solution (including DMSO), without the APAP and the CF™ fluorescent dyes.

12 segments of 3 *µ*L each were produced to cover regularly a logarithmic scale between the low and high concentration solutions. An additional 3 *µ*L segment of the control solution was added before partitioning. These segments were injected in a droplet production chip with the following geometrical parameters: an injector width and height respectively of 100 and 40 *µ*m and a slope of 8 %. The volume of the first (with the cells) and second (with the drug) droplets was respectively estimated to 12 nL and 60 nL, so the dilution factor from the library to the post-merging droplets was approximately 6.

### Droplet Library Injection in the Anchors Array

During the segment partitioning, the droplets were kept in the storage chamber of the microfluidic droplet producing chip. Before injection into a trapping chamber with capillary anchors, the droplet producing chip was manually flipped over several times to mix the different droplet types inside the storage chamber Fig. S4*C*. Then, the chip was connected to the aqueous inlet of anchor array chip, and it was maintained upside down to allow the droplets to escape the storage chamber Fig. S4*E*. Using the aqueous inlet for the droplet injection presented several advantages. The separation between the droplets in the trapping chip was controlled by the oil flowrate at the junction inlet and it also broke down possible large droplets coming from a coalescence event during the droplet transfer. In addition, the flowrate in the chamber was controlled independently with the chamber inlet.

### Image Acquisition

Images without cells were acquired either on a binocular (MZ16 FA, Leica) using a CCD camera (Insight camera, 4MP Firewire, SPOT), or, for the colored droplets, with a digital single-lens reflex camera (D7000, Nikon). The droplet production images of Fig. S3*B* were acquired with a high speed camera (FAST- CAM 1024 PCI, Photron). The cellular fluorescent images were taken with an inverted microscope (Eclispe Ti, Nikon), equipped of a motorized stage, an illumination system (Spectra-X, Lumencor) and a temperature controlled incubator (Okolab), with a CMOS camera (ORCA Flash 4.0, Hamamatsu).

### Image Analysis

For quantifying the droplet colors in Fig. 2, the entire array was imaged using the binocular and the reflex camera. Then, a custom Matlab code (R2016a, Mathworks) allowed to detect each anchor and to compute the RGB values in the center of each droplet, before and after merging.

For the toxicity experiment, single images of the anchors were acquired automatically with the motorized stage of the microscope. The analysis was conducted on a montage of the detected anchors using a protocol previously described (21). Briefly, cells were detected using bright field and fluorescent intensities, and spheroids were selected based on morphological parameters. For each spheroid, the local background was used to determine a specific threshold for the fluorescent dead cells. The viability at the spheroid level was then defined as:

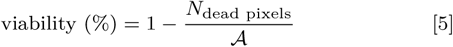

*N*_dead pixels_ and 𝒜 being respectively the number of dead pixels and area of the spheroid. A similar analysis was conducted for determining the viability of the 2D cultures.

At the cellular level, dead cell centers were detected as the local maxima of the fluorescent mortality marker, above the local threshold. Then, the radial distance of each dead cell center was computed and compared to the equivalent radius R of the spheroid to define the mean normalized distance of dead cell centers to the spheroid center 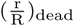. It was close to 0 if all detected dead cells were close to the spheroid center and close to 1 if they were close to the spheroid edge.

### Optical Determination of the Acetaminophen Concentration in Droplets

The drug concentration in the droplets was determined right after the merging of the spheroid and drug droplets by measuring the signals from the CF™ dyes. For each anchor, the fluorescent signal was defined as the local background (average of the signal just outside this anchor) subtracted from the raw fluorescent intensity (average of the raw intensity in the middle of the deep part of the anchor). This signal was correlated to a drug concentration using the calibration curves of the CF™ dyes shown in Fig. S9. In order to replicate the conditions of the experiments, the calibration was made by trapping large droplets of culture medium in the deep parts of the anchors with a similar concentration of viability dyes, DMSO and a known concentration of each CF™ dye. 4 concentrations were loaded in each chip at the same time (21), giving about 60 fluorescent droplets per concentration. 2 chips were used for getting the 8 measurements of each calibration curve. The droplets were imaged with the same experimental parameters than for the toxicity experiments.

### Statistical analysis

*: p < 0.05; **: p < 0.01; ***: p < 0.001; N.S.: non-significant. p-value ranges are only indicated for the highlighted comparisons. Details of each statistical test and p-values can be found in Table S2. In the Tukey box-and-whiskers figures (Fig. 6), the boxes represent the first (q1) and third (q3) quartiles with the median shown by the line bissecting the box, and the mean is shown with black circles. The whiskers represent 1.5 times the inter-quartile range (q3-q1) of the sample. Finally, the box width is proportional to 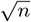.

## ACKNOWLEDGMENTS

Caroline Frot is gratefully acknowledged for her help with the microfabrication and Gabriel Amselem for useful discussions. The A673 cells were kindly provided by Karine Laud Duval from the U830 INSERM/Institut Curie unit. The research leading to these results received funding from the European Research Council (ERC) Grant Agreement 278248 Multicell.

